# Decoupling bile acid 7α-dehydroxylation from colonization resistance to *Clostridioides difficile*

**DOI:** 10.1101/2025.05.20.655077

**Authors:** Luca Beldi, Yuan Dong, Colin Volet, Disha Tandon, Kristýna Filipová, Rizlan Bernier-Latmani, Siegfried Hapfelmeier

**Author notes:** Correspondence to: Siegfried Hapfelmeier. Yuan Dong: genOway Shanghai, Co., Ltd.;, Disha Tandon: Faculty of Science, Institute of Biology, University of Neuchâtel, Switzerland;, Colin Volet: Metabolomics and Proteomics Platform (MAPP), University of Fribourg, Fribourg, Switzerland.

## Abstract

Secondary bile acids, generated through microbial transformation of primary bile acids secreted in bile, play a role in shaping intestinal microbial communities, modulating host immunity, and regulating energy metabolism. In vitro studies have shown that the balance between primary and secondary bile acids strongly affects spore germination, growth, and cellular physiology of *Clostridioides difficile*, a major nosocomial gut pathogen. In vivo correlations between microbiome composition, bile acid metabolome, and colonization resistance have led to the hypothesis that 7α-dehydroxylating bacteria such as *Clostridium scindens* protect against *C. difficile* infection by producing secondary bile acids like deoxycholic acid. However, due to the genetic intractability of known 7α-dehydroxylating species, direct experimental validation of this causal relationship has been challenging.

In this study, we leveraged the first available 7α-dehydroxylation-deficient *baiH* mutant to test the direct role of 7-dehydroxylated bile acid production in *C. difficile* colonization resistance in vivo. We colonized gnotobiotic mice with isogenic wild-type or *baiH* strains of the recently described 7α-dehydroxylating species *Faecalicatena contorta*, including wild-type *C. scindens*-colonized mice as a positive control. Wild-type *F. contorta* accumulated 7-dehydroxylated bile acids at levels equivalent to *C. scindens*, in a strictly *baiH*-dependent manner. However, despite equivalent bile acid profiles, wild-type *F. contorta* failed to replicate the *C. difficile*-restrictive phenotype observed with *C. scindens*.

These findings demonstrate that commensal clostridial 7α-dehydroxylation alone is not sufficient for enhancing colonization resistance to *C. difficile*. Our results highlight the existence of additional, potentially bile acid-independent mechanisms by which certain commensals mediate protection, with important implications for microbiota-based therapies.

**Importance:** 7α-dehydroxylated secondary bile acids, including deoxycholic acid and lithocholic acid, produced by commensal clostridia are widely assumed to inhibit the important nosocomial pathogen *Clostridioides difficile*, yet their precise role in colonization resistance remains unresolved. Using a defined mouse microbiota and an isogenic *Faecalicatena contorta* strain pair differing in a single 7α-dehydroxylation gene (*baiH*), we show that restoration of secondary bile acid production is not sufficient to delay *C. difficile* colonization in vivo. This contrasts with the protective effect of *Clostridium scindens*, which generates a similar bile acid profile. Our findings uncouple bile acid metabolism from protection and suggest that additional, strain-specific functions – such as nutrient competition or antimicrobial production – play a critical role. Understanding these mechanisms is essential for the rational design of next-generation microbiota-based therapies to prevent or treat recurrent *C. difficile* infection.

## Introduction

Bile acids are cholesterol-derived molecules synthesized in the liver that play essential roles in digestion, primarily by solubilizing dietary lipids and fat-soluble vitamins in the small intestine [1]. In addition to their digestive function, bile acids are increasingly recognized as signalling molecules with immunomodulatory and host metabolic regulatory activity, acting through host receptors such as the Farnesoid X Receptor (FXR) and the G-protein-coupled bile acid receptor 1 (TGR5) [2]. Beyond their direct effects on host physiology, bile acids both influence and are influenced by the gut microbiota, with wide-ranging effects on host health and the gut microbiota-mediated colonization resistance against enteric pathogens such as *Clostridioides difficile* [3–5].

In humans, the primary bile acids cholic acid (CA) and chenodeoxycholic acid (CDCA) are synthesized in the liver, conjugated with either glycine or taurine (with taurine conjugation being almost exclusive in mice [6]), and secreted into the proximal small intestine via the biliary tract [7, 8]. While CA and CDCA are the main primary bile acids in humans, in mice, CA and muricholic acids (αMCA and βMCA) – derivatives of CDCA that are absent in humans – predominate [9]. Approximately 95% of secreted primary bile acids are reabsorbed from the small intestine and recycled via the enterohepatic circulation. The remaining 5% (equivalent to 400-800 mg/day in humans) reach the more densely colonized large intestinal lumen, where they undergo extensive microbial transformation into a diverse array of secondary bile acids. Some of these enzymatic modifications, including taurine/glycine deconjugation, oxidations, and epimerizations, can be carried out by a wide range of gut microbial species [10]. However, the 7α-dehydroxylation of CA and CDCA, yielding the major secondary bile acids deoxycholic acid (DCA) and lithocholic acid (LCA), respectively, is restricted to a limited diversity of species within the Bacillota phylum, specifically members of the *Lachnospiraceae*, *Oscillospiraceae*, *Peptostreptococcaceae*, and *Ruminococcaceae* families [11]. This transformation is mediated by the bile acid-inducible (*bai*) gene operon encoding enzymes catalysing the multi-step 7α-dehydroxylation pathway [12, 13]. The presence of *bai* gene-encoding species in the microbiota has been repeatedly correlated with enhanced colonization resistance against *C. difficile* [5, 14–16].

*Clostridioides difficile* (formerly *Clostridium difficile* [17]) is a Gram-positive, obligate anaerobic, endospore-forming bacterium and a major cause of antibiotic-associated diarrhea in humans [18, 19]. Its ability to form endospores confers high resistance to environmental stressors, including alcohol-based disinfectants, making it a persistent threat in healthcare settings [20–22]. Under normal conditions, the resident gut microbiota provides effective colonization resistance against *C. difficile* by still incompletely understood mechanisms [23]. Antibiotic treatment can disrupt this microbial barrier by altering microbiome composition and function, depleting key taxa, and thereby reducing colonization resistance - ultimately predisposing the host to *C. difficile* infection (CDI) [24, 25].

Bile acids have been shown to play key regulatory roles in the life cycle of *C. difficile*, particularly in vitro [26]. Primary bile acids such as CA and taurocholic acid (TCA) as well as DCA, together with the co-germinant glycine, promote *C. difficile* spore germination [5, 13, 27]. In contrast, CDCA and the secondary bile acids LCA and ursodeoxycholic acid (UDCA) inhibit this process [28]. The vegetative growth of *C. difficile* is also negatively affected by bile acids, likely due to their detergent properties that disrupt bacterial membrane integrity. Among these, 7α-dehydroxylated secondary bile acids DCA and LCA (and derivatives thereof) exhibit the strongest inhibitory effects in a dose dependent manner [29]. In co-culture experiments, 7α-dehydroxylating bacterial species inhibited *C. difficile* specifically in the presence of CA, correlating with the production of DCA in a strain-dependent manner [30]. In vivo studies in mice found strong negative correlations between the abundances of 7α-dehydroxylating species, 7α-dehydroxylated bile acids, and *C. difficile* colonization resistance [14–16, 31].

Among currently characterized 7α-dehydroxylating organisms, *Clostridium scindens* has emerged as particularly effective. Its administration to antibiotic-pretreated conventional mice reduced the initial severity of subsequent *C. difficile* infection, providing the first demonstration of an individual commensal species contributing to *C. difficile* colonization resistance in vivo [15]. This modest but reproducible delay of infection has been confirmed in two independent studies using gnotobiotic mouse models lacking endogenous 7α-dehydroxylation activity – one harbouring a defined 12-member murine microbiota [14], the other a 37-member human-derived community [32]. In mentioned three studies, *C. scindens* alone could not confer full colonization resistance but significantly delay *C. difficile* infection, associated with the partially restored accumulation of 7α-dehydroxylated bile acids. These findings have led to prevailing view that production of 7α-dehydroxylated bile acids is a principal mechanism by which *C. scindens* and related 7α-dehydroxylating species contribute to colonization resistance against *C. difficile* in vivo.

However, the support of this hypothesis has largely relied on in vitro models or aforementioned correlative in vivo data. A recent study using *Cyp8b1* knockout mice, which lack CA and all its derivatives, suggested that *C. scindens* may mediate protection predominantly through bile acid-independent mechanisms, potentially by competing with *C. difficile* for Stickland fermentation substrates such as proline [33]. Additional studies have directly linked proline limitation to *C. difficile* colonization resistance: low intestinal proline levels correlated with reduced *C. difficile* burdens in a human microbiota-associated mouse model [34]. The requirement of proline for growth has long been recognized [35] and recently shown to mediate the competition between *C. difficile* and commensal clostridia consortia or human fecal microbiota during in vitro cultivation [36, 37]. Proline availability is also implicated in the regulation of *C. difficile* virulence [38] and toxin expression [39]. Other host-derived nutrients, including sorbitol [40, 41], sialic acid [42], and N-acetylglucosamine [43], and various amino acids like leucine and ornithine [44] also have been shown to influence *C. difficile* colonization dynamics.

Although *Cyp8b1* knockout mice reportedly exhibit no overt changes in microbiota composition, potential effects arising from the global absence of CA-derived bile acids or compensatory production of alternative bile acid species cannot be entirely excluded [33]. A more direct and controlled approach would isolate the specific contribution of microbial 7α-dehydroxylation activity to colonization resistance – without altering host bile acid biosynthesis or microbiota species composition – through head-to-head comparison of isogenic wild-type and *bai* gene mutant strains in vivo. Until recently, this had not been feasible due to the lack of genetically intractable 7α-dehydroxylating strains. However, the recent development of a targeted *bai* gene mutant strain has enabled such type of experiment [45]. Specifically, a newly isolated strain of *Faecalicatena contorta* (*F. contorta*^WT^), a member of the *Lachnospiraceae* and close phylogenetic relative of *C. scindens*, was genetically modified by disruption of the *baiH* gene, generating a 7α-dehydroxylation-deficient isogenic mutant (*F. contorta*^Mut^) [45].

In this study we hypothesized that colonization of mice with *F. contorta*^WT^ would reproduce the transient colonization resistance against *C. difficile* as observed with *C. scindens* [14], whereas the colonization of *F. contorta*^Mut^ would not impact *C. difficile* colonization, mimicking the phenotype of negative control mice. To test this in a clean and reproducible system, we used gnotobiotic OligoMM^12^ mice that harbour a rationally designed mouse derived 12-species bacterial microbiota devoid of any endogenous 7α-dehydroxylating activity [14, 46]. The gnotobiotic OligoMM^12^ mouse model has emerged as a versatile in vivo platform for precision microbiome analysis, enabling specific microbial augmentation experiments [14, 46–49]. OligoMM^12^ efficiently deconjugates primary bile acids, required for studying microbial 7-dehydroxylation which requires deconjugated bile acids as substrates. This model was augmented with either wild-type *F. contorta*, its isogenic *baiH* mutant, or wild type *C. scindens* as a positive control. We then directly compared their bile acid and microbiota compositions and assessed the resulting levels of colonization resistance against *C. difficile*.

## Results

### *Faecalicatena contorta* colonizes OligoMM^12^ mice without skewing endogenous microbiota composition

We first assessed whether *Faecalicatena contorta* strain S122 (*F. contorta*^WT^) and its isogenic 7α-dehydroxylation-deficient mutant (genotype Ω*baiH*; *F. contorta*^Mut^) [45] could stably colonize OligoMM^12^ mice without disrupting the existing microbiota. Previous studies have shown that the OligoMM^12^ consortium supports modular augmentation with additional strains including *C. scindens* [14, 49–51]. Gnotobiotic mice harbouring the OligoMM^12^ community were orally inoculated with 10^9^ CFU of either *F. contorta*^WT^ or *F. contorta*^Mut^. A third group was colonized with *C. scindens* ATCC35704 wild type, previously shown to partially restore bile acid 7α-dehydroxylation and colonization resistance against *C. difficile* [14] (**Fig 1A**).

**Figure 1.**
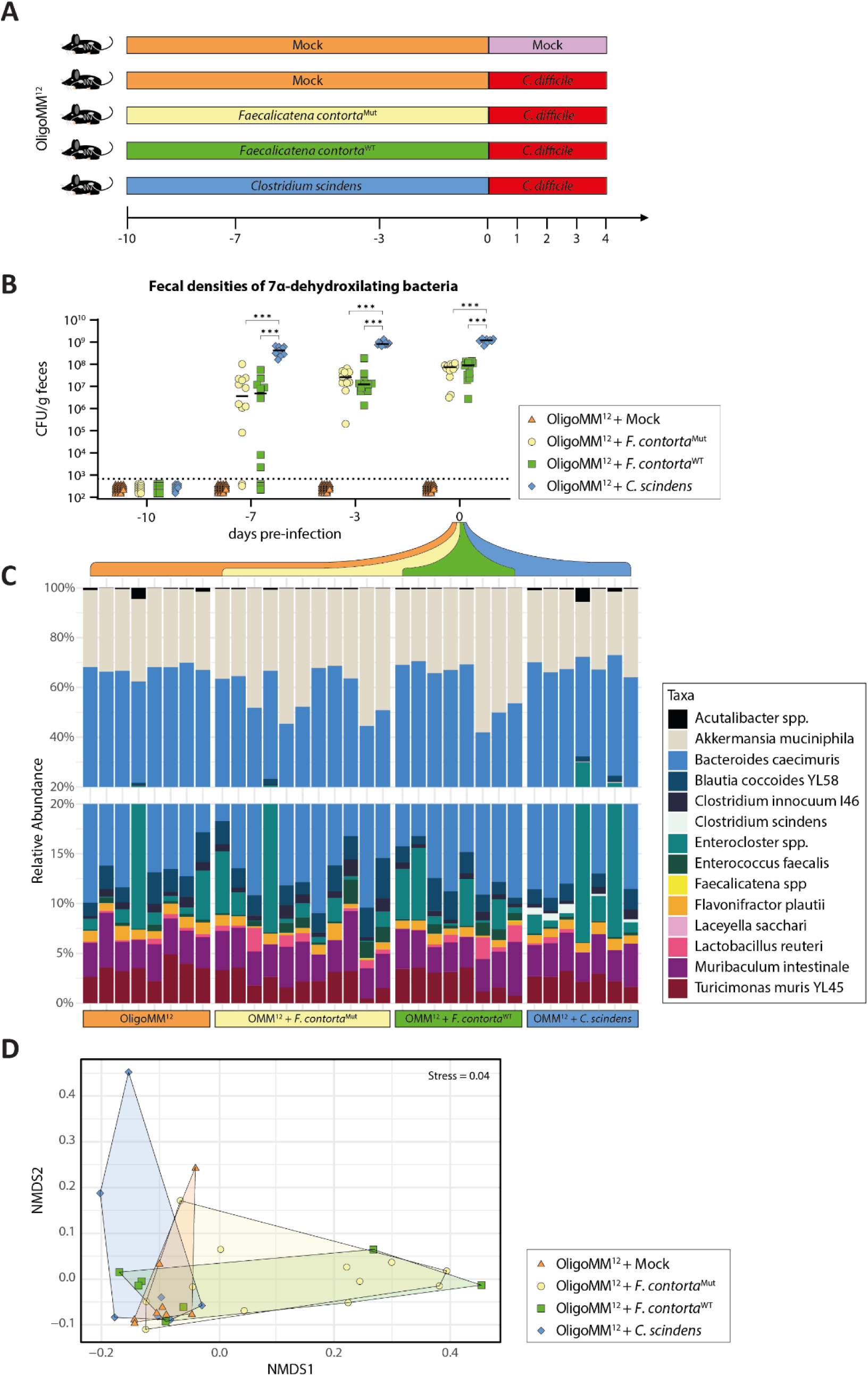
*F. contorta* and *C. scindens* colonization in OligoMM^12^ mice does not change consortium composition. (**A**) C57BL/6 mice colonized with the OligoMM^12^ consortium were recruited into 5 experimental groups and colonized for ten days with either *Faecalicatena contorta*^WT^ (n=12), *F. contorta*^Mut^ (n=12) or *C. scindens* (n=9) or mock colonized with PBS (n=22). At time point 0 mice were orally administered with 10^3^ spores of *C. difficile* or mock infected with PBS and fecal pellets were collected at the indicated time points (-10, -7, -3, 0, 1, 2, 3). Mice were euthanized 4 days post infection. (**B**) Colonization with 7α-dehydroxylating species *F. contorta*^WT^, *F. contorta*^Mut^ and *C. scindens* was confirmed by CFU enumeration on selective plates prior to infection. (**C**) Relative abundances of bacterial species in the colonized mice was evaluated prior to infection (t=0) by compositional analysis of full length 16S rRNA amplicon nanopore-sequencing. (**D**) Non-parametric multidimensional scaling of the relative abundance data shown in (**C**). Each symbol represents one individual. Bars indicate medians. Dotted line marks the lower limit of detection. Statistical analysis in (**B**) used 2-way ANOVA to compare colonization of 7α-dehydroxylating bacteria and excluded mock colonized animals (\*\*\**p*<0.001).

Fecal CFU counts over 10 days confirmed successful colonization of both *F. contorta* strains. Although initial densities varied widely between mice on day -7 (pre-infection), both *F. contorta*^WT^ and *F. contorta*^Mut^ reached similar steady-state levels by the time of infection (10^7.64^ ^± 0.53^ and 10^7.72^ ^± 0.51^ CFU/g feces, respectively; geometric mean × geometric SD factor) (**Fig 1B**). Extended observation beyond 10 days revealed no further increases in population densities (**Fig S1A-B**). Fecal CFU counts of *C. scindens* showed less variability and significantly higher steady-state levels (10^9.06^ ^± 0.10^ CFU/g feces; p < 0.001 vs. *F. contorta* colonized groups; mock-amended OligoMM^12^ controls excluded from statistical comparison). While plating efficiency differences between both species may contribute, 16S rRNA gene amplicon sequencing confirmed these trends (**Fig 1C**). *C. scindens* was detectable in all mice, with a mean relative abundance of 0.45% (±0.28% SD), whereas *F. contorta*^WT^ and *F. contorta*^Mut^ were detected in only 3 out of 19 mice, indicating ≥10-fold lower relative abundances.

Importantly, the addition of either 7α-dehydroxylating strain did not significantly alter the composition of the core OligoMM^12^ community (adjusted p > 0.05, pairwise PERMANOVA). Non-metric multidimensional scaling (NMDS) analysis of relative abundance data showed overlapping clustering across all groups (**Fig 1D**), supporting the conclusion that microbiota structure remained stable following augmentation.

### *baiH*-dependent production of secondary bile acids in *F. contorta*-colonized mice recapitulates the bile acid metabolome of *C. scindens*-colonized mice

Having established stable colonization of both *F. contorta* strains, we next asked whether *F. contorta*^WT^ could functionally restore 7α-dehydroxylation-dependent secondary bile acid metabolism in OligoMM^12^ mice (**Fig 2A, B**). After 10 days of colonization (corresponding to day 0 as defined in **Fig 1A**, prior to *C. difficile* infection), we quantified fecal concentrations of 40 bile acid species using ultra-high-performance liquid chromatography coupled to high-resolution mass spectrometry (UHPLC-HRMS; see Methods). Bile acid 7α-dehydroxylation requires the prior deconjugation of primary bile acids by the OligoMM^12^ community, as neither *F. contorta* nor *C. scindens* encodes bile salt hydrolases required for this step. Consistent with earlier studies [14, 50, 52], the OligoMM^12^ core microbiota supports bile acid de-conjugation and some additional transformations, but lacks bile acid 7α-dehydroxylation activity (**Fig S2** and **Fig 2A**). Colonization with either wild-type *F. contorta* or *C. scindens*, introducing the *bai* operon, restored bile acid 7α-dehydroxylation (**Fig S2** and **Fig 2C**).

**Figure 2.**
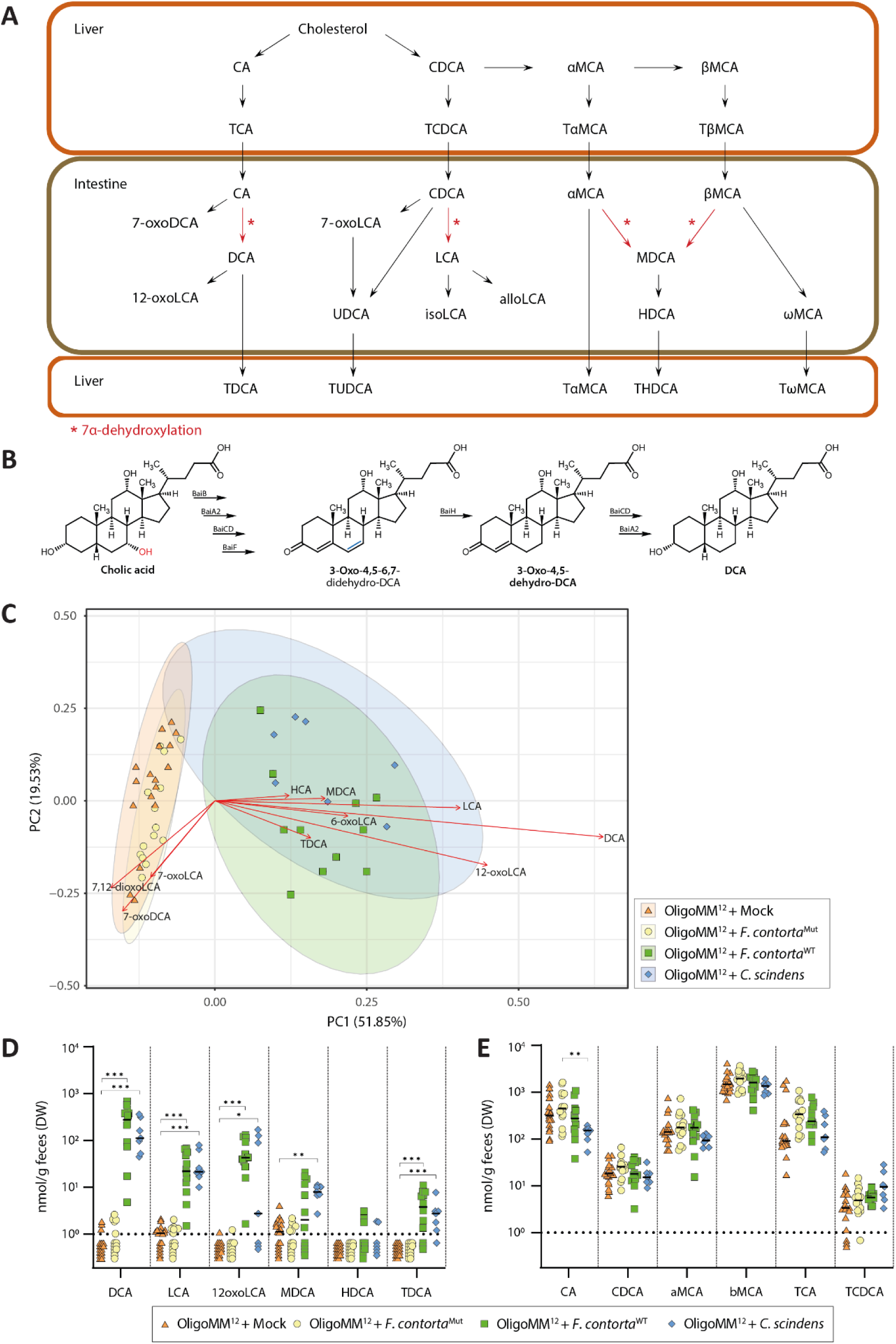
Bile acid metabolomes of OligoMM^12^ mice colonized with *F. contorta*^WT^ or *C. scindens* are indistinguishable. (**A**) Schematic representation of murine bile acid metabolism (adapted from Studer et al 2016 [14]). (**B**) Simplified biosynthetic scheme of bile acid 7α-dehydroxylation (adapted from Jin 2022 [45]). The *baiH* gene encodes for an oxidoreductase responsible for the reduction of 3-Oxo-4,5-6,7-didehydro-DCA to 3-Oxo-4,5-dehydro-DCA. *F. contorta*^Mut^ was shown to accumulate the predicted intermediate and fails to produce DCA [45]. (**C**) Principal component analysis of fecal bile acid metabolites concentrations quantified before *C. difficile* infection (day 0). Red arrows represent the top 10 loadings of principal component 1. Ellipses indicate 95% confidence intervals. (**D, E**) Fecal concentrations of (**D**) 7α-dehydroxylated bile acids, (**E**) deconjugated primary bile acids, and the main *C. difficile* germinants TCA and TCDCA prior to *C. difficile* colonization. Statistical analysis in panels D and E was performed using Kruskal-Wallis multiple comparison test with Dunn’s post hoc test. Each symbol represents one individual. Bars indicate median values. Dotted line marks lower limit of quantification (1nmol/g feces). (\**p*<0.05, \*\**p*<0.01, \*\*\**p*<0.001). 12-oxoLCA, 12-oxolithocholic acid; 6-oxoLCA, 6-oxolithocholic acid; 7,12-dioxoLCA, 7,12-dioxolithocholic acid; 7-oxoDCA, 7-oxodeoxycholic acid; 7-oxoLCA, 7-oxolithocholic acid; alloLCA, allo-lithocholic acid; αMCA, α-murocholic acid; βMCA, β-murocholic acid; CA, cholic acid; CDCA, chenodeoxcholic acid; DCA, deoxycholic acid; isoLCA, iso-lithocholic acid; HCA, hyocholic acid; HDCA, hyodeoxycholic acid; LCA, lithocholic acid; MDCA, murodeoxycholic acid; ωMCA, ω-muricholic acid; TαMCA, tauro-α-murocholic acid; TβMCA, tauro-β-murocholic acid; TCA, taurocholic acid; TCDCA, taurochenodeoxycholic acid; TDCA, taurodeoxycholic acid; THDCA, taurohyodeoxycholic acid; TωMCA, tauro-ω-muricholic acid; TUDCA, tauroursodeoxycholic acid; UDCA, ursodeoxycholic acid.

Unsupervised hierarchical clustering of bile acid profiles revealed two distinct groups: mice colonized with *F. contorta*^WT^ or *C. scindens* clustered together, while those colonized with 7α-dehydroxylation-deficient *F. contorta*^Mut^ and mock-treated OligoMM^12^ controls formed a separate cluster (**Fig S2**). Principal component analysis (PCA) analysis identified the 7α-dehydroxylated bile acids DCA, LCA, and MDCA, along with their derivatives, as the major drivers of variation along PC1, explaining the observed clustering (**Fig 2C**). Notably, there was no separation between wild type *F. contorta*^WT^*-* and *C. scindens-*colonized mice, indicating that both organisms generated similar large intestinal secondary bile acid profiles.

Non-parametric multiple comparisons of 7α-dehydroxylation dependent secondary bile acids confirmed this similarity: concentrations of DCA and LCA – two secondary bile acids implicated in suppressing vegetative *C. difficile* colonization [14, 29] – and other 7α-dehydroxylated bile acids did not differ significantly between *F. contorta*^WT^- and *C. scindens*-colonized mice (**Fig 2D**). In contrast, mice colonized with *F. contorta*^Mut^ or mock controls exhibited only trace or undetectable levels of these bile acids.

Deconjugated primary bile acids CA, CDCA, αMCA, and βMCA – the educts of 7α-dehydroxylation –showed no significant differences between groups, except for CA, which was present at statistically significantly different levels between *C. scindens-* and *F. contorta*^Mut^-colonized mice (**Fig 2E**). Concentrations of the conjugated primary bile acids taurocholic acid (TCA) and taurochenodeoxycholic acid (TCDCA), known to promote *C. difficile* spore germination [5, 13, 27], did not vary significantly between groups, suggesting that bile acid-mediated spore germination cues were not substantially altered.

### *C. scindens-*mediated delay of *C. difficile* colonization is not phenocopied by *F. contorta*^WT^

We next investigated whether 7α-dehydroxylation activity alone was sufficient to confer colonization resistance against *C. difficile.* Comparing *F. contorta*^WT^ with its isogenic *baiH* mutant (*F. contorta*^Mut^) in OligoMM^12^ mice enabled gene-level testing of 7α-dehydroxylation function in mediating *C. difficile* colonization resistance. On day 10 of pre-colonization, in parallel with fecal bile acid profiling (see **Figure 2**), mice from each group (*F. contorta*^WT^-, *F. contorta*^Mut^-, *C. scindens*- or mock*-*colonized) were challenged with *C. difficile* endospores. Mock-infected OligoMM^12^ mice served as uninfected controls.

To compare their colonization resistance, fecal *C. difficile* CFU densities were monitored daily over 4 days (**Fig 3A-B**). Consistent with prior work [14, 15], pre-colonization with *C. scindens* significantly delayed *C. difficile* infection. In contrast, mice colonized by *F. contorta*^WT^ showed no such delay and were indistinguishable from those colonized with the 7-dehydroxylation-deficient *F. contorta*^Mut^ strain (**Fig 3A-B**). *C. difficile* colonization dynamics in these groups also mirrored those of mock-colonized (OligoMM^12^ community-only) controls. *C. difficile-*induced colonic inflammation, assessed by quantitation of cecal luminal lipocalin-2 [14], followed the same trend (**Fig 3C**).

**Figure 3.**
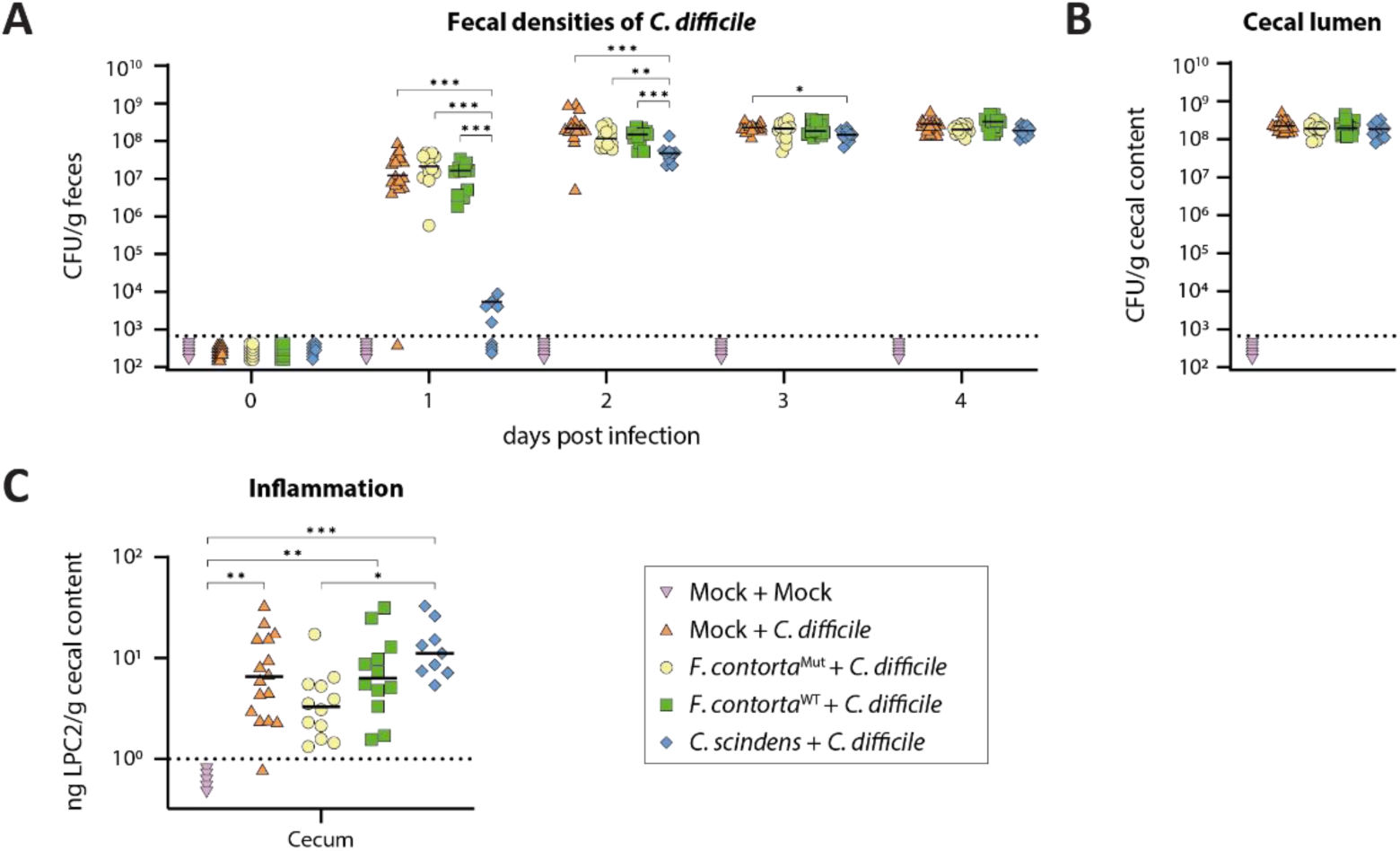
Delayed colonization phenotype of *C. difficile* is only observed in *C. scindens* colonized OligoMM^12^ mice. (**A**) All mice were infected with *C. difficile* spores or mock infected with PBS on day 0. Fecal pellets were collected on days 1-4. Fecal densities of *C. difficile* CFU were assessed by selective plating. (**B**) Cecal *C. difficile* CFU numbers were terminally assessed on day 4 by selective plating. (**C**) Measured lipocalin-2 (LPC2) in ng/g feces of collected cecum samples on day 4 post infection. Points represent individual mice. Dotted lines represent the lower limit of detection. Statistical analysis passed Shapiro-Wilk lognormality test (excluding mock infected animals) and compared groups with a 2-way Anova and Tukey’s multiple comparisons test in (**A&B**) and used Kruskal Wallis test with Dunn’s multiple comparison test in (**C**). (* *p* < 0.05, ** *p* < 0.01, *** *p* < 0.001).

Together, these findings demonstrate that *F. contorta* recapitulates intestinal bile acid 7α-dehydroxylation but not the *C. difficile-*suppressing effects of *C. scindens*. This suggests that bile acid 7α-dehydroxylation alone is insufficient to enhance colonisation resistance and that *C. scindens* likely employs additional mechanisms to suppress *C. difficile* in vivo.

## Discussion

In this study, we demonstrate that although wild-type *Faecalicatena contorta* can stably colonize OligoMM^12^ mice and reconstitute 7α-dehydroxylation-dependent secondary bile acid production, this metabolic activity alone is not sufficient to delay *C. difficile* colonization. This contrasts with the protective effect observed in *C. scindens*-colonized mice, which exhibited nearly identical fecal bile acid profiles, and suggests that bile acid 7α-dehydroxylation alone cannot fully account for the protective phenotype. Our results highlight the importance of considering additional strain-specific functions – beyond bile acid metabolism – that may contribute to colonization resistance.

We first confirmed that wild-type *F. contorta* stably colonized OligoMM^12^ mice without disrupting the existing intestinal community and reached steady-state densities similar to its isogenic *baiH*-deficient mutant. Although *F. contorta* achieved lower fecal abundances than *C. scindens*, targeted metabolomic profiling revealed that it restored a nearly identical secondary bile acid profile, including key secondary bile acids such as DCA and LCA. The comparison with a *baiH* targeted *F. contorta* mutant provided a genetically controlled system to dissect the contribution of bile acid 7α-dehydroxylation in vivo.

Despite this bile acid metabolic restoration, wild type *F. contorta* failed to delay *C. difficile* colonization relative to its *baiH* mutant or to mock-colonized controls. These results show that bile acid 7α-dehydroxylation alone does not enhance colonization resistance in vivo and indicate that other functional traits of *C. scindens* are likely responsible for its protective effect. Such traits may include competitive exclusion or direct antagonism through secretion of other bioactive molecules.

This interpretation is supported by previous studies reporting bile acid-independent mechanisms of *C. scindens*-mediated inhibition. For example, coculture assays revealed that some *C. scindens* strains suppress *C. difficile* even when bile acid transformation is limited, suggesting additional inhibitory mechanisms beyond 7α-dehydroxylation [30]. Kang et al. identified antimicrobial tryptophan-derived metabolites produced by strains of *C. scindens* (and *C. sordellii*) that synergize with secondary bile acids to inhibit *C. difficile* in vitro [26]. More recently, *C. scindens-*secreted molecules distinct from bile acids were reported to downregulate *C. difficile* virulence secretion [53]. Additionally, work in *Cyp8b1*-deficient mice (lacking CA production) mono-colonized with *C. scindens* revealed that the individual protective potential of *C. scindens* against *C. difficile* in vivo is most likely mediated not by bile acids but by competition for Stickland fermentation substrates, particularly proline [33]. Competition for proline has emerged as a dominant determinant of *C. difficile* colonization resistance, contributed also by non-bile acid transforming organisms [32, 36, 37]. In this study, we did not investigate the energy metabolism or antimicrobial biosynthetic potential of *F. contorta*, but we hypothesise that it lacks bile acid-independent protective traits comparable to those of *C. scindens*.

By decoupling bile acid metabolism from *C. difficile* suppression in a genetically fully defined in vivo system – including host genetic background, a fully defined microbiota, and manipulation of a single *bai* gene – we provide a more precise framework for evaluating microbial functions that contribute to colonization resistance. Our findings emphasize the importance of assessing the functional context and microbial background in which bacterial metabolic pathways operate.

In conclusion, while secondary bile acids remain an important component of the host-microbial crosstalk regulating gut microbiota and host physiology, their protective role against *C. difficile* colonization is context-dependent and likely synergizes with other, strain-specific microbial traits. Identifying and harnessing such traits – particularly those employed by commensals like *C. scindens* – will be essential for the rational design of defined, standardized therapeutic microbial consortia.

## Materials & methods

### Animals and ethics statement

Animal experiments were conducted with age matched (7-13Weeks), gender-balanced and littermate-controlled C57BL/6J mice. Mice were bred under strict gnotobiotic conditions in the Clean Mouse Facility (EAC University of Bern) with tightly controlled housing conditions. The ambient temperature was between 23-25°C with a relative humidity between 52-60%. Mice received autoclaved rodent chow diet Kliba-Nafag 3307 (Granovit) and autoclaved surgical irrigation water (Baxter Healthcare ERKF7114) *ad libitum* at all times.

Gnotobiotic OligoMM^12^ mice have been derived from germ-free C57BL/6 mice through colonization with the OligoMM^12^ microbiota strains in the Clean Mouse Facility of the University of Bern. These animals were maintained in flexible film isolators before they were exported into individually ventilated cages (IVC, model Sealsafe Plus, Tecniplast) for colonization with *F. contorta* and *C. scindens*. OligoMM^12^ mice are stably colonized with 12 strains (*Acutalibacter muris* KB18 (DSM 26090), *Akkermansia muciniphila* YL44 (DSM 26127), *Bacteroides caecimuris* I48 (DSM 26085), *Bifidobacterium longum* subsp. *animalis* YL2 (DSM 26074), *Blautia pseudococcoides* YL58 (DSM 26115), *Clostridium innocuum* I46 (DSM26113), *Enterocloster clostridioformis* YL32 (DSM 26114), *Enterococcus faecalis* KB1 (DSM 32036), *Flavonifractor plautii* YL31 (DSM26117), *Limosilactobacillus reuteri* I49 (DSM 32035), *Muribaculum intestinale* YL27 (DSM 28989), *Turicimonas muris* YL45 (DSM 26109)) all of which are available at https://www.dsmz.de/collection/catalogue/microorganisms/special-groups-of-organisms/dzif-sammlung/maus-mikrobiomliste. OligoMM^12^ microbiota stability and reproducibility *in vivo* have been addressed in great detail previously [46, 54].

*C. difficile* infections were performed either in the Central Animal Facility (CAF, Institute for Pathology) or in a clean mouse room (CMR, Institute for infectious disease) at the University of Bern. For the duration of the infection, mice were housed in individually ventilated cages (IVC, model Sealsafe Plus, Tecniplast).

All animal experiments were approved by the Bernese Cantonal ethical committee for animal experiments under the license number BE66/19 and BE44/22 and were in accordance with the Swiss Federal law and the Bernese Cantonal regulations.

### Bacterial culture conditions and CFU quantitation

*C. scindens* ATCC35704 [55] and *Faecalicatena contorta* S122 (both WT and Ω*baiH* mutant) [45] were grown in *C. scindens* medium containing 37 g Brain Heart Infusion (BD 237500), 5 g yeast extract (BD 212750), 1 g L-cystein (Merck 2838), 2 g Fructose (Merck 5321) and 40 ml *C. scindens* Salts Solution (0.2 g CaCl_2_ (Merck 2083), 0.2 g MgSO_4_ (Merck 5886), 1 g K_2_HPO_4_ (Merck 5101), 1 g KH_2_PO_4_ (Merck 4873), 10 g NaHCO_3_ (Merck 6329), and 2 g NaCl (Merck 6404) in 1 L _dd_H_2_O) per 1 liter _dd_H_2_O. Erythromycin (20 µg/ml, Merck E5389-5G) and Nalidixic acid (50 µg/ml, Grogg N8878) were added to *C. scindens* medium agar plates for monitoring colonization prior to *C. difficile* infection. Plates were imported in an anaerobic workstation (Don Whitley A45 HEPA, 10% CO_2_, 10% H_2_, 80% N_2_) 24 h prior to usage allowing residual oxygen to diffuse out of the agar.

*Clostridioides difficile* DH1916 [56] was grown in BHIY containing 37 g Brain Heart Infusion (BD 237500), 5 g yeast extract (BD 212750), 0.3 g L-cysteine (Merck 2838) per 1 Liter _dd_H_2_O. For selective *C. difficile* colony forming unit enumeration we used *C. difficile* Fructose Agar which contains 69 g *C. difficile* Agar Base (Fluka 17145), 5 g yeast extract (Oxoid LP21), 250 mg D-cycloserine (Carl Roth CN37.3), 8 mg Cefoxitin (Sigma C4786) and 1 g Taurocholate (Sigma 86339) per 1 Liter _dd_H_2_O. Plates were imported in an anaerobic workstation (Don Whitley A45 HEPA, 10% CO_2_, 10% H_2_, 80% N_2_) 24 h prior to usage allowing residual oxygen to diffuse out of the agar.

Collected intestinal contents (feces, colonic and cecal contents) were weighed and resuspended in 1 ml pre-reduced PBS for 3 min in a tissue Lyser (LT, Qiagen) at 50 HZ. Fecal suspension was plated on selective plates and incubated for either 1 day (*C. difficile*) or 3 days (7α-dehydroxylating bacteria) at 36°C prior to CFU enumeration.

### Pre-colonization with 7*α*-dehydroxylating bacteria

OligoMM^12^ mice were exported from their breeding isolator into IVC cages and administered orally with 10^9^ CFU (in a volume of 200 µl) of either *C. scindens* or *F. contorta*. Control animals were administered orally with 200 µl PBS (80 g sodium chloride (Merck 106404), 2 g potassium chloride (Merck 104936), 14.4 g disodium hydrogen phosphate (Merck 106586), 2.4 g potassium dihydrogen phosphate (Merck 104873) per 1 Liter _dd_H_2_O). Pre-colonization periods lasted for 10 days and regular faecal sampling confirmed colonization with the respective applied strain.

### Preparation of *C. difficile* spores and mouse infections

*C. difficile* DH1916 spores were prepared by growing 4x 10 ml cultures in 70:30 sporulation medium [57] for 5 days. Subsequently, cultures were frozen at -20°C for 2 days. Spores were washed at least 10 times with ice-cold _dd_H_2_O until spore purity reached >95%. Spores were heated at 65°C for 30 min and then stored at 4°C in PBS+1% BSA upon infection of animals. Spore preparation purity was checked by phase contrast microscopy and was enumerated through plating on BHIY plates containing 0.1% taurocholate.

Mice were infected with 10^3^ CFU of spores in a total volume of 200 µl PBS. Fecal pellets (24 h, 48 h, 72 h and 96 h post-infection) or cecal contents (24 h or 96 h post-infection after sacrifice) were collected for *C. difficile* enumeration on selective plates.

### Bile acid analysis

Samples were lyophilized overnight at -60°C. The dried fecal contents were weighed and 6 ceramic beads (2.5 mm) were added to each tube. 750 µl of MeOH/H2O (2/1) + 0.1% formic acid was used as extraction solvent. Samples were homogenised in a Precellys 24 Tissue Homogenizer (Bertin Instruments) at 6500 rpm 2x 20 sec beat and 20 sec rest. The homogenized cecal samples were centrifuged at 21’000 *g*, for 15 min, at 4°C. 100 µl from each supernatant or calibration standard were transferred into individual wells of 2 ml 96-well plate. 50 µl of an ISTD solution (CA-d4, CDCA-d4, TCA-d4, TUDCA-d4, DCA-d4 and LCA-d4, each at 2 μM in methanol) was pipetted in each well. Immediately after the addition of ISTD, 600 µl of 0.2% formic acid in H_2_O was added to each sample or calibration standard level. The 96-well plate was shaken with an orbital shaker at 300 rpm and centrifuged at 3500 rpm for 5min at 4°C. The contents of the 96-well plate were extracted by solid phase extraction with an Oasis HLB 96-well uElution plate. The extracted samples were dried in a Biotage® SPE Dry 96 at 20°C and reconstituted with 100 µl of MeOH/H_2_O (50/50). The plate was shaken with an orbital shaker at 300 rpm for 5min and centrifuged at 3500 rpm for 5 min at 4°C. The samples were injected on the LC-HRMS system.

The quantitative method was performed on a Thermo Vanquish Flex High performance liquid chromatography coupled in tandem to a Thermo Exploris 120 mass spectrometer [58]. The separation was done on a Zorbax Eclipse Plus C18 column (2.1 × 100 mm, 1.8 μm) and a guard column Zorbax Eclipse Plus C18 (2.1 × 5 mm, 1.8 μm) both provided by Agilent technologies. The column compartment was kept heated at 50°C. Two different solutions were used as eluents: ammonium acetate [5 mM] in water as mobile phase A and pure acetonitrile as mobile phase B. A constant flow of 0.4 ml/min was maintained over 26 minutes of run time with the following gradient (expressed in eluent B percentage): 0-5.5 min, constant 21.5% B; 5.5-6 min, 21.5-24.5% B; 6-10 min, 24.5-25% B; 10-10.5 min, 25-29% B; 10.5-14.5 min, isocratic 29% B; 14.5-15 min, 29-40% B; 15-18 min, 40-45% B; 18-20.5 min, 45-95% B; 20.5-23 min, constant 95% B; 23-23.1 min, 95-21.5% B; 23.10-26 min, isocratic 21.50% B. The system equilibration was implemented at the end of the gradient for 3 minutes in initial conditions. The autosampler temperature was maintained at 10°C and the injection volume was 5 μl. The ionisation mode was operated in negative mode for the detection using H-ESI. The Exploris 120 acquisition settings were configured in resolution mode 120000 FWHM at m/z 200, data storage in profile and centroid mode and the high-resolution full MS chromatograms were acquired over the range of m/z 200-700. The mass spectrometer was calibrated in negative mode using ESI calibrant solution from Thermo to maintain the best possible mass accuracy.

Data were processed afterwards using the Thermo TraceFinder Quantitative software and Thermo Freestyle Qualitative software to control the mass accuracy for each run. In the quantitative method, 43 bile acids were quantified by calibration curves. The quantification was corrected by addition of internal standards in all samples and calibration levels. Extracted ion chromatograms were generated using a retention time window of ± 1.5 min and a mass extraction window of ± 10 ppm around the theoretical mass of the targeted bile acid.

### Lipocalin-2 quantification

Lipocalin-2 quantification was performed similar as previously described [14, 49] using a commercially available mouse lipocalin-2/NGAL ELISA DuoSet (R&D DY1857). The wells of a 96- well plate were coated with lipocalin-2 capture antibody (1 ng/well in 50 µl PBS, incubated at 4°C over-night), washed with a non-ionic detergent washing buffer (0.05% Tween-20 (Merck 93773) in PBS) and blocked with blocking buffer (PBS with 2% bovine serum albumine (Merck 10735078001), 150 µl per well for 3 h at RT). Diluted fecal and cecal samples (10-fold dilutions in PBS, 4 times) and standard (3-fold dilutions in PBS, 8 times in duplicate) were dispensed onto the wells (50 µl per well) and incubated for 3 h at RT. After washing, 50 µl of biotinylated anti-lipoclain-2 antibody (25 ng/well in blocking buffer) was incubated for 1 h at RT. Another washing step was performed before adding 100 µl of streptavidin conjugated horseradish-peroxidase (40-fold diluted in PBS) for 1 h at RT. Wells were washed again with washing buffer prior to development using 100 µl substrate buffer (0.1M NaH_2_PO_4_ (Merck 1063451000), pH 4) with 0.01% ABTS (Merck 11557) and 0.1% H_2_O_2_ (Merck 216763-100ML). The absorbance was measured at 405 nm with a plate reader (THERMOmax microplate reader). Measured values were fitted using non-linear regression and lipocalin-2 concentrations of samples were back-calculated to ng lipocalin-2 per gram sample.

### 16S rRNA gene amplicon sequencing analysis

Intestinal content samples from mice were collected in a tube and resuspended in sterile PBS. Metagenomic DNA was extracted as per the instructions in the kit PureLink™ Microbiome DNA Purification Kit (Invitrogen A29790). Quality control and quantification of the genomic DNA was done using Nanodrop One^C^ (ND-ONEC-W Thermo Fisher Scientific) 260/230 and 260/280 ratios. The concentration of all DNA samples was determined by Qubit dsDNA BR kit (Life Technologies Q32853) on the Qubit 4.0 fluorometer.

PCR amplification of the 16S rRNA gene was performed using modified degenerate universal primers 27Fdeg-1492Rdeg (**Table 1**). PCR conditions include initial denaturation at 95° C for 5 minutes, followed by 30 thermal cycles of denaturation at 94° C for 30 seconds, annealing at 58° C for 30 seconds and elongation at 72° C for 40 seconds. This was followed by final end product extension at 72° C for 10 minutes. PCR was followed by gel electrophoresis to confirm the amplification. PCR amplicons were purified using Minelute PCR purification kit (Qiagen 28006). PCR purified amplicons were end-prepared using Native Barcoding Kit 24 V14 (Oxford Nanopore Technologies SQK-NBD114.24) according to manufacturer’s instruction with minor modifications. The end-prepared amplicons were purified using AMPure XP beads (Beckman Coulter SQK-NBD114.24) on the DynaMAG.96 Side skirted Magnetic plate (Thermo Fisher Scientific 12027). The purified amplicons were barcoded using the native kit, according to manufacturer’s instructions, except increased starting amounts of amplicon DNA (250 ng/amplicon). The barcoded amplicon DNA was pooled in equal concentrations and purified using AMPure XP beads on the DynaMAG.96 Side skirted Magnetic plate. In the final step, adapter ligation was carried out using Ligation Sequencing Kit V14 (Oxford Nanopore Technologies SQK-LSK114) according to manufacturer’s instructions. Sequencing runs were performed on the R10.4.1 flow cell (Oxford Nanopore Technologies FLO-MIN114).

**Table 1.**
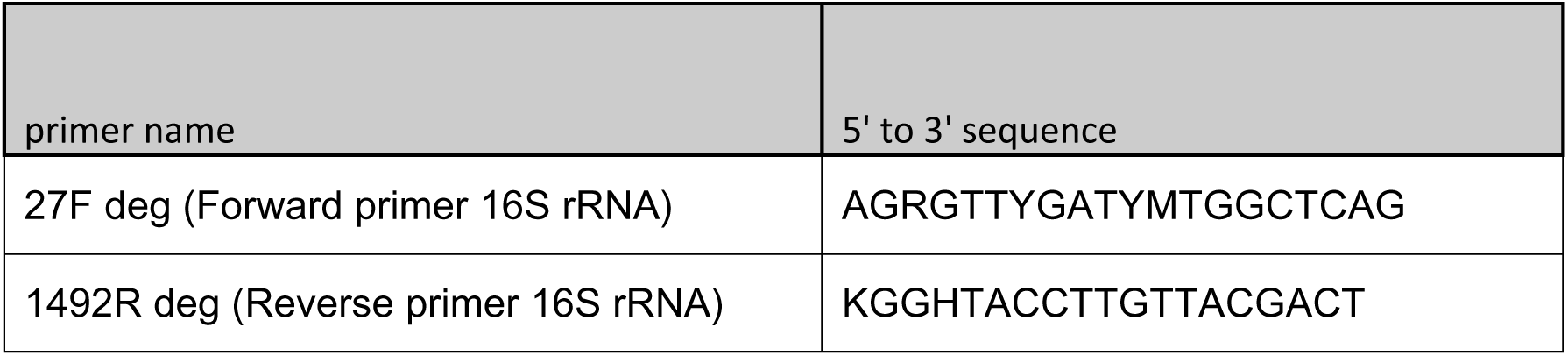
Primer sequences.

**Table 2.** Bacterial strains.

| Bacterial Strain | Genetic background | Relevant genotype | Antibiotic resistances | Relevant phenotype | Reference |
| --- | --- | --- | --- | --- | --- |
| *Clostridioides difficile* DH1916 | Ribotype 027 | Wild type | cefoxitin, D-cycloserine, erythromycin |  | [56] |
| *Faecalicatena contorta*<sup>WT</sup> | *Faecalicatena contorta* S122 | Wild type | nalidixic acid, erythromycin, D-cycloserine, |  | [45] |
| *Faecalicatena contorta*<sup>Mut</sup> | *Faecalicatena contorta* S122 | *ΩbaiH* | nalidixic acid, erythromycin, D-cycloserine, thiamphenicol | 7α-dehydroxylation-deficient | [45] |
| *Clostridium scindens* ATCC35704 | ATCC35704 | Wild type | erythromycin |  | [55] |

Samples with more than 1000 reads were considered for further analysis. Raw reads were base-called and demultiplexed using the ’guppy’ (version:5.1.13+b292f4d13, default mode) [59]. Chimeras were checked by mapping the reads (all-against-all) using ’minimap2’ [60] and then detecting the chimeric reads using ’yacrd’ [61]. The reads were filtered by quality (>=9) and length (1000-1600) using ’chopper’ [62]. Reads were aligned and classified using ’EMU’ [63] against a custom database that includes sequences from rrnDB v5.6 [64] and NCBI 16S RefSeq [65] databases.

### Statistical analyses

Statistical analysis was performed as described in Figure legends using Graph Pad Prism V10.2.1. or RStudio (Version 2024.12.1+563). Heatmap hierarchical clustering was done using the dist function (“canberra”) and hclust (“average”) in R. PCA used the prcomp function and for the NMDS the distance matrix was created using the function vegdist (“bray” method) and results were generated with the metaMDS function.

### Software

ChatGPT4.0 was utilized to correct language and style

### Data availability

Raw reads of 16S data are available in the public repository ENA under the accession number PRJEB89465. Other data is available in supplementary source_data_file.

## Acknowledgments

This work was funded by SNF Sinergia grant #CRSII5_180317 and SNF NCCR Microbiomes grants #51NF40_180575 (Phase I) and #51NF40_ 225148 (Phase II). The funding bodies had no role in study design, data collection and interpretation, or the decision to submit the work for publication. Animal experimental support and infrastructure were provided by the central animal facility (CAF) and the clean mouse facility (CMF) of the University of Bern. We thank Chun-Jun Guo (University of Cornell) for providing *Faecalicatena contorta* strains, Alban Ramette (IFIK, University of Bern) for providing statistical consultation, Olivier Schären, Matheus Notter Dias and all Hapfelmeier lab members for scientific discussions and experimental support.

## Author contributions

Conceptualization: LB, YD, SH; Data curation: LB, YD, CV, DT; Formal analysis: LB, YD; Funding acquisition: SH; Investigation: LB, YD, CV, KF, DT; Methodology: LB, YD, SH; Project administration: LB, YD, SH; Resources: SH, RBL; Supervision: SH; Validation: LB, SH; Visualization: LB; Writing-original draft: LB, SH; Writing-review & editing: All authors

---

**Figure S1.**
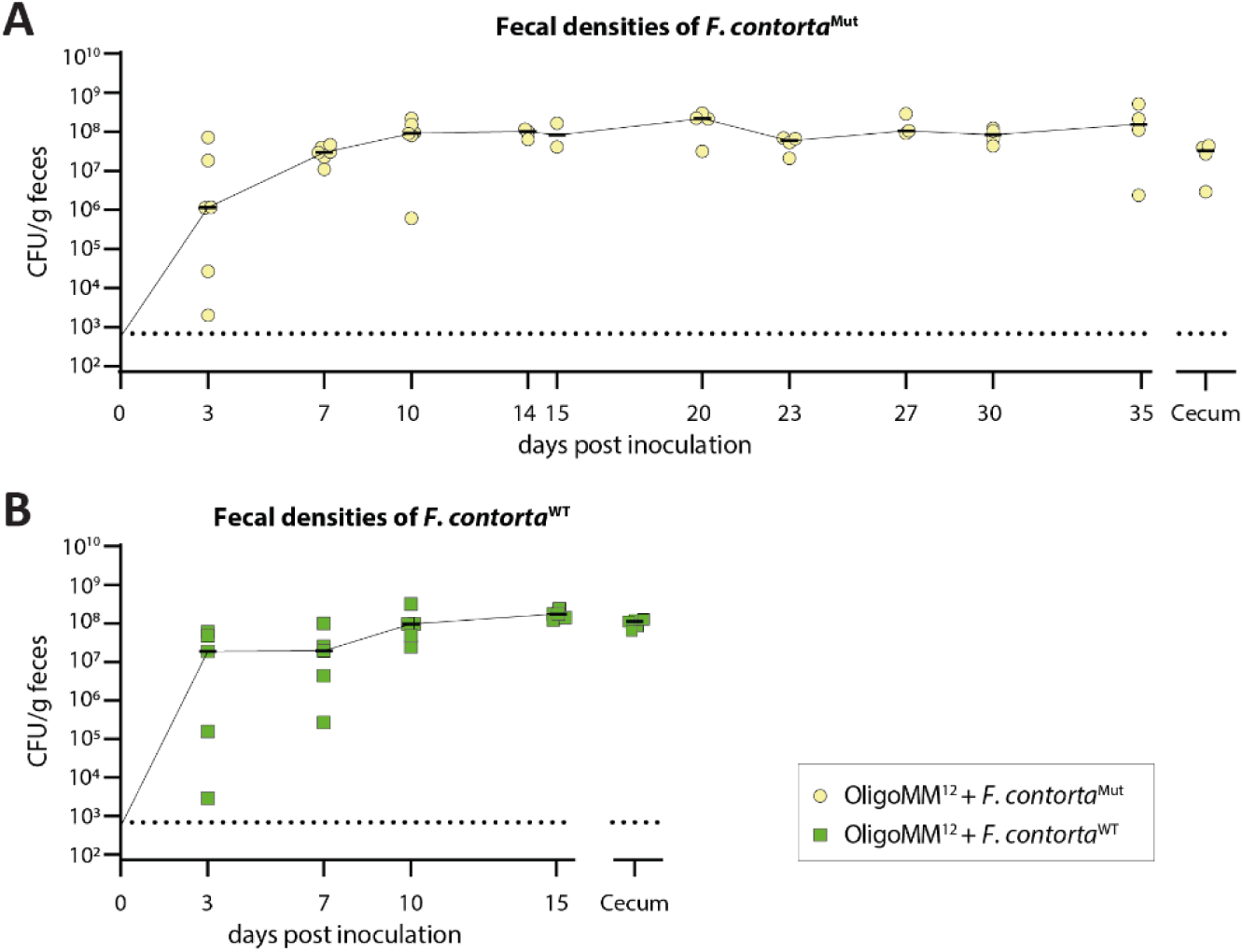
Extended colonization of *F. contorta* in OligoMM^12^ mice. OligoMM^12^ mice were inoculated with either *F. contorta*^Mut^ (**A**) or *F. contorta*^WT^ (**B**) and sampled at indicated time points. CFU/g feces were evaluated by selective plating. Dotted lines represent lower limits of detection. Bars indicate medians. For CFU/g in cecum quantification, content was collected at experimental endpoint and also plated on selective plates.

**Figure S2.**
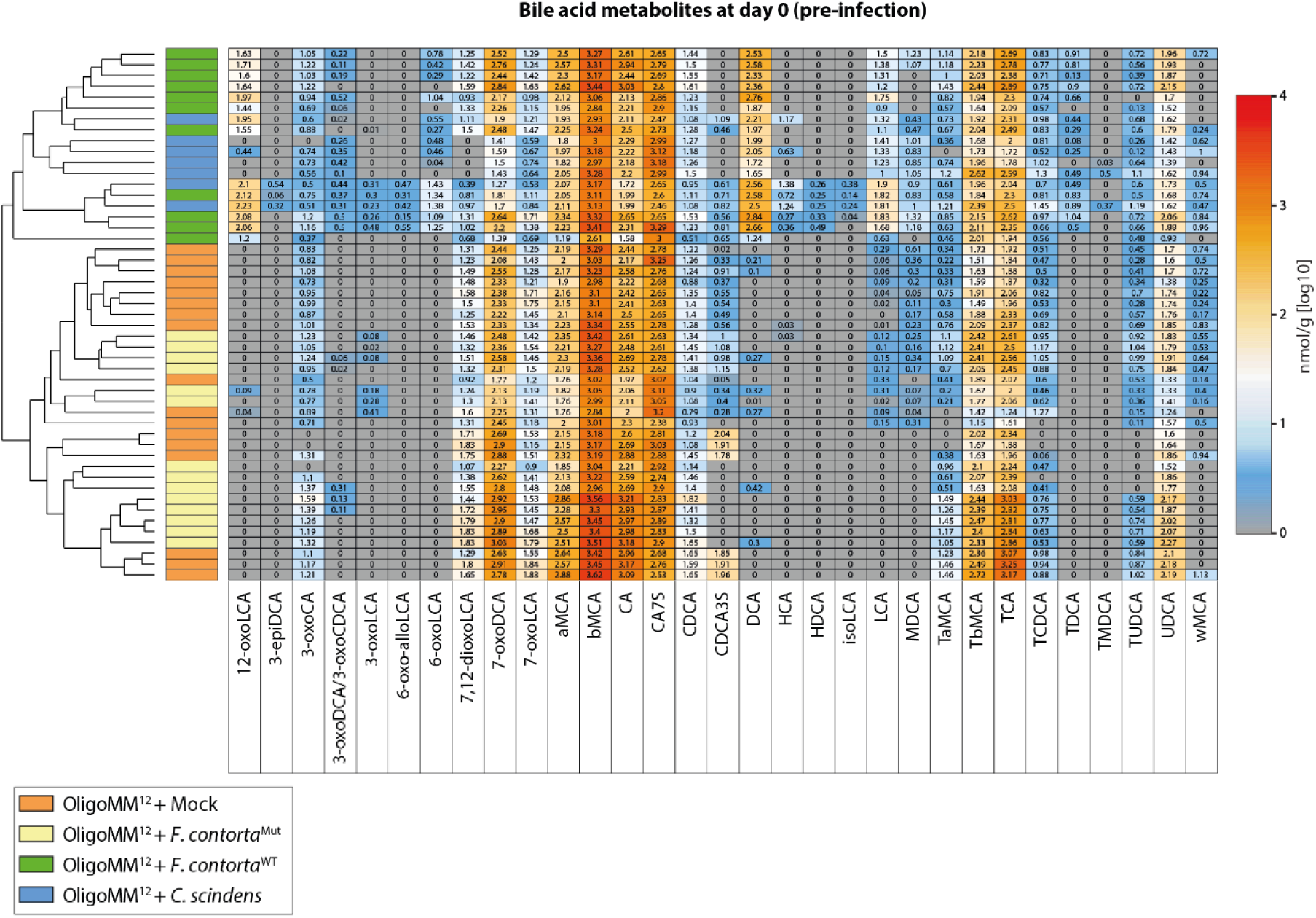
Unsupervised hierarchical clustered heatmap from measured bile acid compounds. Values represent measured bile acid concentrations in nmol/g (log10) dry weight of fecal content measured via UHPLC-HRMS. Samples were collected at time point of infection (t=0). 12-oxoLCA, 3-epiDCA, 3-epideoxycholic acid; 3-oxoCA, 3-oxocholic acid; 3-oxoDCA, 3-oxodeoxycholic acid; 3-oxoCDCA, 3-oxochenodeoxycholic acid; 6-oxo-alloLCA, 6-oxo-allolithocholic acid; 6-oxoLCA, 6-oxolithocholic acid; 7,12-dioxoLCA, 7,12-dioxolithocholic acid; 7-oxoDCA, 7-oxodeoxycholic acid; 7-oxoLCA, 7-oxolithocholic acid; αMCA, α-murocholic acid; βMCA, β-murocholic acid; CA, cholic acid; CA7S, cholic acid 7-sulfate; CDCA, chenodeoxcholic acid; CDCA3S, chenodeoxycholic acid 3-sulfate; DCA, deoxycholic acid; HCA, hyocholic acid; HDCA, hyodeoxycholic acid; isoLCA, iso-lithocholic acid; LCA, lithocholic acid; MDCA, murodeoxycholic acid; TαMCA, tauro-α-murocholic acid; TβMCA, tauro-β-murocholic acid; TCA, taurocholic acid; TCDCA, taurochenodeoxycholic acid; TDCA, taurodeoxycholic acid; TMDCA, tauromurideoxycholic acid; TUDCA, tauroursodeoxycholic acid; UDCA, ursodeoxycholic acid; ωMCA, ω-muricholic acid.

## References

1. Hofmann, A.F., The continuing importance of bile acids in liver and intestinal disease. Arch Intern Med, 1999. 159(22): p. 2647–58 DOI: 10.1001/archinte.159.22.2647.

2. Fiorucci, S., et al., Bile Acids Activated Receptors Regulate Innate Immunity. Front Immunol, 2018. 9: p. 1853 DOI: 10.3389/fimmu.2018.01853.

3. Larabi, A.B., H.L.P. Masson, and A.J. Bäumler, Bile acids as modulators of gut microbiota composition and function. Gut Microbes, 2023. 15(1) DOI: Artn 2172671 10.1080/19490976.2023.2172671.

4. Ramirez-Perez, O., et al., The Role of the Gut Microbiota in Bile Acid Metabolism. Ann Hepatol, 2017. 16 Suppl 1: p. S21–S26 DOI: 10.5604/01.3001.0010.5672.

5. Winston, J.A. and C.M. Theriot, Impact of microbial derived secondary bile acids on colonization resistance against Clostridium difficile in the gastrointestinal tract. Anaerobe, 2016. 41: p. 44–50 DOI: 10.1016/j.anaerobe.2016.05.003.

6. Falany, C.N., et al., Cloning, expression, and chromosomal localization of mouse liver bile acid CoA:amino acid N-acyltransferase. J Lipid Res, 1997. 38(6): p. 1139–48.

7. Falany, C.N., et al., Glycine and taurine conjugation of bile acids by a single enzyme. Molecular cloning and expression of human liver bile acid CoA:amino acid N-acyltransferase. J Biol Chem, 1994. 269(30): p. 19375–9.

8. Vessey, D.A., The biochemical basis for the conjugation of bile acids with either glycine or taurine. Biochem J, 1978. 174(2): p. 621–6 DOI: 10.1042/bj1740621.

9. Wahlstrom, A., et al., Intestinal Crosstalk between Bile Acids and Microbiota and Its Impact on Host Metabolism. Cell Metabolism, 2016. 24(1): p. 41–50 DOI: 10.1016/j.cmet.2016.05.005.

10. Ridlon, J.M., D.J. Kang, and P.B. Hylemon, Bile salt biotransformations by human intestinal bacteria. J Lipid Res, 2006. 47(2): p. 241–59 DOI: 10.1194/jlr.R500013-JLR200.

11. Ridlon, J.M. and H.R. Gaskins, Another renaissance for bile acid gastrointestinal microbiology. Nat Rev Gastroenterol Hepatol, 2024. 21(5): p. 348–364 DOI: 10.1038/s41575-024-00896-2.

12. Wells, J.E. and P.B. Hylemon, Identification and characterization of a bile acid 7alpha-dehydroxylation operon in Clostridium sp. strain TO-931, a highly active 7alpha-dehydroxylating strain isolated from human feces. Appl Environ Microbiol, 2000. 66(3): p. 1107–13 DOI: 10.1128/AEM.66.3.1107-1113.2000.

13. Reed, A.D. and C.M. Theriot, Contribution of Inhibitory Metabolites and Competition for Nutrients to Colonization Resistance against Clostridioides difficile by Commensal Clostridium. Microorganisms, 2021. 9(2) DOI: 10.3390/microorganisms9020371.

14. Studer, N., et al., Functional Intestinal Bile Acid 7alpha-Dehydroxylation by Clostridium scindens Associated with Protection from Clostridium difficile Infection in a Gnotobiotic Mouse Model. Front Cell Infect Microbiol, 2016. 6: p. 191 DOI: 10.3389/fcimb.2016.00191.

15. Buffie, C.G., et al., Precision microbiome reconstitution restores bile acid mediated resistance to Clostridium difficile. Nature, 2015. 517(7533): p. 205–8 DOI: 10.1038/nature13828.

16. Solbach, P., et al., BaiCD gene cluster abundance is negatively correlated with Clostridium difficile infection. PLoS One, 2018. 13(5): p. e0196977 DOI: 10.1371/journal.pone.0196977.

17. Lawson, P.A., et al., Reclassification of Clostridium difficile as Clostridioides difficile (Hall and O’Toole 1935) Prevot 1938. Anaerobe, 2016. 40: p. 95–99 DOI: 10.1016/j.anaerobe.2016.06.008.

18. Zhu, D., J.A. Sorg, and X. Sun, Clostridioides difficile Biology: Sporulation, Germination, and Corresponding Therapies for C. difficile Infection. Front Cell Infect Microbiol, 2018. 8: p. 29 DOI: 10.3389/fcimb.2018.00029.

19. Sebaihia, M., et al., The multidrug-resistant human pathogen Clostridium difficile has a highly mobile, mosaic genome. Nat Genet, 2006. 38(7): p. 779–86 DOI: 10.1038/ng1830.

20. Jabbar, U., et al., Effectiveness of alcohol-based hand rubs for removal of Clostridium difficile spores from hands. Infect Control Hosp Epidemiol, 2010. 31(6): p. 565–70 DOI: 10.1086/652772.

21. Bouza, E., Consequences of Clostridium difficile infection: understanding the healthcare burden. Clin Microbiol Infect, 2012. 18 Suppl 6: p. 5–12 DOI: 10.1111/1469-0691.12064.

22. Lemiech-Mirowska, E., et al., The Hospital Environment as a Potential Source for Clostridioides difficile Transmission Based on Spore Detection Surveys Conducted at Paediatric Oncology and Gastroenterology Units. Int J Environ Res Public Health, 2023. 20(2) DOI: 10.3390/ijerph20021590.

23. Pike, C.M. and C.M. Theriot, Mechanisms of Colonization Resistance Against Clostridioides difficile. J Infect Dis, 2021. 223(12 Suppl 2): p. S194–S200 DOI: 10.1093/infdis/jiaa408.

24. Theriot, C.M., et al., Antibiotic-induced shifts in the mouse gut microbiome and metabolome increase susceptibility to Clostridium difficile infection. Nat Commun, 2014. 5: p. 3114 DOI: 10.1038/ncomms4114.

25. Bartlett, J.G., Antibiotic-associated pseudomembranous colitis. Rev Infect Dis, 1979. 1(3): p. 530–9 DOI: 10.1093/clinids/1.3.530.

26. Kang, J.D., et al., Bile Acid 7alpha-Dehydroxylating Gut Bacteria Secrete Antibiotics that Inhibit Clostridium difficile: Role of Secondary Bile Acids. Cell Chem Biol, 2019. 26(1): p. 27–34 e4 DOI: 10.1016/j.chembiol.2018.10.003.

27. Sorg, J.A. and A.L. Sonenshein, Bile salts and glycine as cogerminants for Clostridium difficile spores. J Bacteriol, 2008. 190(7): p. 2505–12 DOI: 10.1128/JB.01765-07.

28. Sorg, J.A. and A.L. Sonenshein, Inhibiting the initiation of Clostridium difficile spore germination using analogs of chenodeoxycholic acid, a bile acid. J Bacteriol, 2010. 192(19): p. 4983–90 DOI: 10.1128/JB.00610-10.

29. Thanissery, R., J.A. Winston, and C.M. Theriot, Inhibition of spore germination, growth, and toxin activity of clinically relevant C. difficile strains by gut microbiota derived secondary bile acids. Anaerobe, 2017. 45: p. 86–100 DOI: 10.1016/j.anaerobe.2017.03.004.

30. Reed, A.D., et al., Strain-Dependent Inhibition of Clostridioides difficile by Commensal Clostridia Carrying the Bile Acid-Inducible (bai) Operon. J Bacteriol, 2020. 202(11) DOI: 10.1128/JB.00039-20.

31. Theriot, C.M., A.A. Bowman, and V.B. Young, Antibiotic-Induced Alterations of the Gut Microbiota Alter Secondary Bile Acid Production and Allow for Clostridium difficile Spore Germination and Outgrowth in the Large Intestine. mSphere, 2016. 1(1) DOI: 10.1128/mSphere.00045-15.

32. Tian, S., et al., A designed synthetic microbiota provides insight to community function in Clostridioides difficile resistance. Cell Host Microbe, 2025. 33(3): p. 373–387 e9 DOI: 10.1016/j.chom.2025.02.007.

33. Aguirre, A.M., et al., Bile acid-independent protection against Clostridioides difficile infection. PLoS Pathog, 2021. 17(10): p. e1010015 DOI: 10.1371/journal.ppat.1010015.

34. Battaglioli, E.J., et al., Clostridioides difficile uses amino acids associated with gut microbial dysbiosis in a subset of patients with diarrhea. Sci Transl Med, 2018. 10(464) DOI: 10.1126/scitranslmed.aam7019.

35. Karasawa, T., et al., A defined growth medium for Clostridium difficile. Microbiology (Reading), 1995. 141 (Pt 2): p. 371–5 DOI: 10.1099/13500872-141-2-371.

36. Lopez, C.A., et al., Clostridioides difficile proline fermentation in response to commensal clostridia. Anaerobe, 2020. 63: p. 102210 DOI: 10.1016/j.anaerobe.2020.102210.

37. Huang, X., et al., Clostridioides difficile colonization is not mediated by bile salts and utilizes Stickland fermentation of proline in an in vitro model. mSphere, 2025. 10(2): p. e0104924 DOI: 10.1128/msphere.01049-24.

38. Jenior, M.L., et al., Novel Drivers of Virulence in Clostridioides difficile Identified via Context-Specific Metabolic Network Analysis. mSystems, 2021. 6(5): p. e0091921 DOI: 10.1128/mSystems.00919-21.

39. Johnstone, M.A. and W.T. Self, d-Proline Reductase Underlies Proline-Dependent Growth of Clostridioides difficile. J Bacteriol, 2022. 204(8): p. e0022922 DOI: 10.1128/jb.00229-22.

40. Pruss, K.M. and J.L. Sonnenburg, C. difficile exploits a host metabolite produced during toxin-mediated disease. Nature, 2021. 593(7858): p. 261–265 DOI: 10.1038/s41586-021-03502-6.

41. Yang, Z., et al., Host Sorbitol and Bacterial Sorbitol Utilization Promote Clostridioides difficile Infection in Inflammatory Bowel Disease. Gastroenterology, 2023. 164(7): p. 1189–1201 e13 DOI: 10.1053/j.gastro.2023.02.046.

42. Ng, K.M., et al., Microbiota-liberated host sugars facilitate post-antibiotic expansion of enteric pathogens. Nature, 2013. 502(7469): p. 96–9 DOI: 10.1038/nature12503.

43. Pereira, F.C., et al., Rational design of a microbial consortium of mucosal sugar utilizers reduces Clostridiodes difficile colonization. Nat Commun, 2020. 11(1): p. 5104 DOI: 10.1038/s41467-020-18928-1.

44. Smith, A.B., et al., Enterococci enhance Clostridioides difficile pathogenesis. Nature, 2022. 611(7937): p. 780–786 DOI: 10.1038/s41586-022-05438-x.

45. Jin, W.B., et al., Genetic manipulation of gut microbes enables single-gene interrogation in a complex microbiome. Cell, 2022. 185(3): p. 547–562 e22 DOI: 10.1016/j.cell.2021.12.035.

46. Brugiroux, S., et al., Genome-guided design of a defined mouse microbiota that confers colonization resistance against Salmonella enterica serovar Typhimurium. Nat Microbiol, 2016. 2: p. 16215 DOI: 10.1038/nmicrobiol.2016.215.

47. Hoces, D., et al., Metabolic reconstitution of germ-free mice by a gnotobiotic microbiota varies over the circadian cycle. PLoS Biol, 2022. 20(9): p. e3001743 DOI: 10.1371/journal.pbio.3001743.

48. Gul, E., et al., Differences in carbon metabolic capacity fuel co-existence and plasmid transfer between Salmonella strains in the mouse gut. Cell Host Microbe, 2023. 31(7): p. 1140–1153 e3 DOI: 10.1016/j.chom.2023.05.029.

49. Pfister, S.P., et al., Uncoupling of invasive bacterial mucosal immunogenicity from pathogenicity. Nat Commun, 2020. 11(1): p. 1978 DOI: 10.1038/s41467-020-15891-9.

50. Streidl, T., et al., The gut bacterium Extibacter muris produces secondary bile acids and influences liver physiology in gnotobiotic mice. Gut Microbes, 2021. 13(1): p. 1–21 DOI: 10.1080/19490976.2020.1854008.

51. Afrizal, A., et al., Enhanced cultured diversity of the mouse gut microbiota enables custom-made synthetic communities. Cell Host Microbe, 2022. 30(11): p. 1630–1645 e25 DOI: 10.1016/j.chom.2022.09.011.

52. Marion, S., et al., Biogeography of microbial bile acid transformations along the murine gut. J Lipid Res, 2020. 61(11): p. 1450–1463 DOI: 10.1194/jlr.RA120001021.

53. Saenz, C., et al., Clostridium scindens secretome suppresses virulence gene expression of Clostridioides difficile in a bile acid-independent manner. Microbiol Spectr, 2023. 11(5): p. e0393322 DOI: 10.1128/spectrum.03933-22.

54. Eberl, C., et al., Reproducible Colonization of Germ-Free Mice With the Oligo-Mouse-Microbiota in Different Animal Facilities. Front Microbiol, 2019. 10: p. 2999 DOI: 10.3389/fmicb.2019.02999.

55. Morris, G.N., et al., Clostridium Scindens Sp-Nov, a Human Intestinal Bacterium with Desmolytic Activity on Corticoids. International Journal of Systematic Bacteriology, 1985. 35(4): p. 478–481 DOI: Doi 10.1099/00207713-35-4-478.

56. Burns, D.A., et al., Reconsidering the sporulation characteristics of hypervirulent Clostridium difficile BI/NAP1/027. PLoS One, 2011. 6(9): p. e24894 DOI: 10.1371/journal.pone.0024894.

57. Edwards, A.N. and S.M. McBride, Isolating and Purifying Clostridium difficile Spores. Methods Mol Biol, 2016. 1476: p. 117–28 DOI: 10.1007/978-1-4939-6361-4_9.

58. Vico-Oton, E., et al., Strain-dependent induction of primary bile acid 7-dehydroxylation by cholic acid. BMC Microbiol, 2024. 24(1): p. 286 DOI: 10.1186/s12866-024-03433-y.

59. Wick, R.R., L.M. Judd, and K.E. Holt, Performance of neural network basecalling tools for Oxford Nanopore sequencing. Genome Biol, 2019. 20(1): p. 129 DOI: 10.1186/s13059-019-1727-y.

60. Li, H., Minimap2: pairwise alignment for nucleotide sequences. Bioinformatics, 2018. 34(18): p. 3094–3100 DOI: 10.1093/bioinformatics/bty191.

61. Marijon, P., R. Chikhi, and J.S. Varre, yacrd and fpa: upstream tools for long-read genome assembly. Bioinformatics, 2020. 36(12): p. 3894–3896 DOI: 10.1093/bioinformatics/btaa262.

62. De Coster, W. and R. Rademakers, NanoPack2: population-scale evaluation of long-read sequencing data. Bioinformatics, 2023. 39(5) DOI: 10.1093/bioinformatics/btad311.

63. Curry, K.D., et al., Emu: species-level microbial community profiling of full-length 16S rRNA Oxford Nanopore sequencing data. Nat Methods, 2022. 19(7): p. 845–853 DOI: 10.1038/s41592-022-01520-4.

64. Stoddard, S.F., et al., rrnDB: improved tools for interpreting rRNA gene abundance in bacteria and archaea and a new foundation for future development. Nucleic Acids Res, 2015. 43(Database issue): p. D593–8 DOI: 10.1093/nar/gku1201.

65. O’Leary, N.A., et al., Reference sequence (RefSeq) database at NCBI: current status, taxonomic expansion, and functional annotation. Nucleic Acids Res, 2016. 44(D1): p. D733–45 DOI: 10.1093/nar/gkv1189.

